# Origin of modern syphilis and emergence of a contemporary pandemic cluster

**DOI:** 10.1101/051037

**Authors:** Natasha Arora, Verena J. Schuenemann, Günter Jäger, Alexander Peltzer, Alexander Seitz, Alexander Herbig, Michal Strouhal, Linda Grillová, Leonor Sánchez-Busó, Denise Kühnert, Kirsten I. Bos, Leyla Rivero Davis, Lenka Mikalová, Sylvia Bruisten, Peter Komericki, Patrick French, Paul R. Grant, María A. Pando, Lucía Gallo Vaulet, Marcelo Rodríguez Fermepin, Antonio Martinez, Arturo Centurion Lara, Lorenzo Giacani, Steven J. Norris, David Šmajs, Philipp P. Bosshard, Fernando González-Candelas, Kay Nieselt, Johannes Krause, Homayoun C. Bagheri

**Author notes:** Current address: Department of Archaeogenetics, Max Planck Institute for the Science of Human History, Jena, Germany (Alexander Herbig, Kirsten I. Bos, Johannes Krause). Current address: Department of Infectious Disease Epidemiology, Imperial College London, UK. Current address: Repsol Technology Center, Madrid, Spain (Homayoun C. Bagheri). Corresponding authors. (N.A.);, (K.N.); (J.K.); (H.C.B.).

## Abstract

Syphilis swept across the world in the 16th century as one of most prominent documented pandemics and is re-emerging worldwide despite the availability of effective antibiotics. Little is known about the genetic patterns in current infections or the evolutionary origins of the disease due to the non-cultivable and clonal nature of the causative bacterium *Treponema pallidum* subsp. *pallidum*. In this study, we used DNA capture and next generation sequencing to obtain whole genome data from syphilis patient specimens and from treponemes propagated in laboratory settings. Phylogenetic analyses indicate that the syphilis strains examined here share a common ancestor after the 15th century. Moreover, most contemporary strains are azithromycin resistant and members of a globally dominant cluster named here as SS14-Ω. This cluster diversified from a common ancestor in the mid-20^th^ century and has the population genetic and epidemiological features indicative of the emergence of a pandemic strain cluster.

## Text

The abrupt onslaught of the syphilis pandemic starting in the late 15^th^ century established this devastating infectious disease as one of the most feared in human history. The first reported outbreaks in Europe were during the War of Naples in 1495 (Quétel, 1990). Subsequently, the epidemic spread to other continents, remaining a severe health burden until the discovery of penicillin in the 20^th^ century. This multistage bacterial infection causes systemic damage through dissemination of the etiological agent *Treponema pallidum* subsp. *pallidum* (TPA). Surprisingly, the last few decades have witnessed a global re-emergence, with number of cases estimated at 10.6 million in 2008 (Rowley et al., 2012). This resurgence despite the availability of effective antibiotics is striking in high-income western nations such as Switzerland, the United Kingdom, and the United States (Fenton et al., 2008; Stamm, 2010). Furthermore, while resistance to penicillin has not been identified in the bacterium, an increasing number of strains fail to respond to the second-line antibiotic azithromycin (Stamm, 2010).

The epidemiological characteristics of this re-emergence are poorly understood, particularly the underlying patterns of genetic diversity. Obtaining genetic data for TPA is hindered by the low quantities of endogenous DNA in clinical samples, and inability to cultivate the pathogen (Fraser et al., 1998). Consequently, much of our molecular understanding comes from propagating strains in laboratory animals to obtain sufficient DNA. The few published whole genomes were obtained after amplification through rabbit passage (L. Giacani et al., 2014; Šmajs, Norris, & Weinstock, 2012; Štaudová et al., 2014), and represent limited diversity for phylogenetic analyses. These sequences suggest that the TPA genome of 1.14 Mb is genetically monomorphic. Potential genetic diversity remains unexplored because clinical samples are mostly typed by PCR amplification of only 1-5 loci from roughly 1000 genes (Grillová et al., 2014; Christina M. Marra et al., 2010). These epidemiological strain typing studies are motivated by the limitations of serologic or microscopic tests to distinguish among TPA strains or among the subspecies *Treponema pallidum* subsp. *pertenue* (TPE) and *Treponema pallidum* subsp. *endemicum* (TEN), which cause the diseases yaws and bejel, respectively. All three diseases are transmitted through skin contact and show an overlap in their clinical manifestations. While syphilis is geographically widespread and generally transmitted sexually, yaws and bejel are mainly found endemically in hot climates and primarily transmitted between children by skin contact (L. Giacani & Lukehart, 2014). The precise relationships among the bacteria are still debated, particularly regarding the evolutionary origin of syphilis.

The paucity of molecular studies and the focus on typing of a few genes means that we have limited information regarding the evolution and spread of epidemic TPA. In this study, we interrogated genome-wide variation across geographically widespread isolates. In total, we obtained 70 samples from 13 countries, including 52 syphilis swabs collected directly from patients between 2012 and 2013, and 18 syphilis, yaws, and bejel samples collected from 1912 onwards and propagated in laboratory rabbits. Through comparative genome analyses and phylogenetic reconstruction, we shed light on the evolutionary history of TPA and identify epidemiologically relevant haplotypes.

### The distinct evolutionary histories of treponemal lineages

Due to the large background of host DNA, extracts from all 70 samples were converted into double stranded Illumina sequencing libraries, then enriched for treponemal DNA using array hybridization capture, and sequenced (supplementary materials) (Hodges et al., 2009; Meyer & Kircher, 2010). The resultant reads (ranging from 483,450 to 100,414,614) were mapped to the TPA reference genome (NIC_REF, NC_021490). Genomic coverage ranged from 0.13-fold to over 1000-fold, with the highest mean coverage in strains propagated in rabbits, and highest variation in the samples collected directly from patients (0.13-fold to 223-fold). We restricted analyses to the 28 samples where at least 80% of the genome was covered by a minimum of three reads. We investigated structural variation through *de novo* assembly for the four highest covered syphilis swab samples (NE17, NE20, CZ27, AU15) and one Indonesian yaws isolate (IND1). Gaps were expected for the 8% of the genome containing repetitive regions and related genes such as the *tpr* subfamilies or the ribosomal RNA operons. We found no significant structural changes in the five genomes (Fig.1A), except for the deletion in IND1 of gene TP1030, which potentially encodes a virulence-factor (Centurion-Lara et al., 2013). The deletion was shared across all the yaws infection isolates, consistent with other studies (Mikalova et al., 2010).

**Fig. 1.**
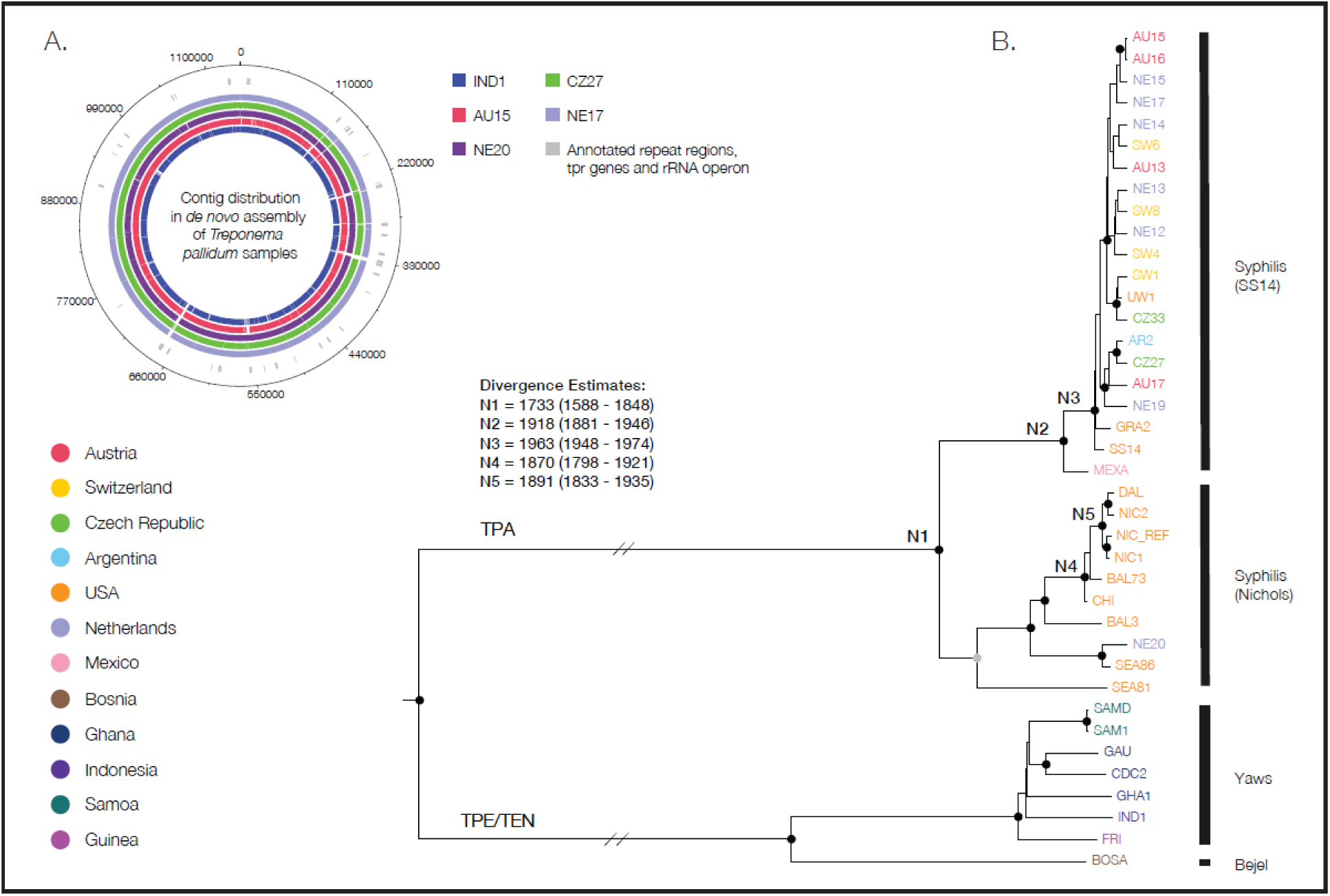
De novo genome assemblies and phylogenetic reconstruction. A) *De novo* genome assembly for four syphilis patient samples and one yaws strain, with color coded geographic origin (inset legend). Blank spaces correspond to gaps, overlapping with gene regions that are difficult to assemble from short reads such as the *tpr* subfamilies and rRNA operons (regions shown in the outermost ring in gray). B) BEAST tree for the 39 genomes (excluding putative recombinant genes), with black circles indicating nodes with ≥99% posterior probabilities (PP); gray circle denotes 82% PP. Divergence date estimates (median and 95% highest posterior density) for major well-supported TPA nodes are given in the legend.

Prior to phylogenetic reconstruction, we checked for signatures of recombination. While *T pallidum* is considered to be a clonal species (Achtman, 2008), previous studies suggest recombinant genes in a Mexican syphilis and a Bosnian bejel strain (Petrosova et al., 2012; Štaudová et al., 2014). We examined each of the 978 annotated genes in our 28 sequenced genomes and the 11 publicly available genomes from laboratory strains. Genes were considered putative recombinants if they satisfied 3 conditions: i) had twice the expected SNP density as compared to the average distribution, ii) produced gene tree topologies incongruent with that of the genome-wide tree, and iii) had four or more homoplasies in at least 2 branches. This procedure identified 4 genes coding for outer membrane proteins, one of which (TP0136) is used in typing studies (Šmajs et al., 2012).

### The independent history and origin of TPA

After removing the 4 putative recombinant genes from the genome alignment (n=2,235 bp) of all 39 genomes, we reconstructed the phylogeny using the Bayesian framework implemented in BEAST (Bouckaert et al., 2014). As illustrated in Fig. 1B, the phylogenetic tree revealed a marked separation of TPA from TPE/TEN, with TPA forming a monophyletic lineage. The distinction of the two lineages was robust even with the inclusion of the putative recombinant genes. Analyses of divergence between the two lineages yielded an average mean distance of 1225 nucleotide differences. By contrast, within each of the lineages we found less than 1/5 the divergence (124.6 average pairwise mutations within the TPA lineage and 200.2 within TPE/TEN). These results highlight the perils of relying on a limited set of markers for taxonomic classification, as they may yield spurious groupings. For instance, phylogenies produced with the typing gene TP0548 do not separate TPA and TPE.

Using the sample isolation dates as tip calibration and applying the Birth Death Serial Skyline model (Stadler, Kuhnert, Bonhoeffer, & Drummond, 2013), we obtained a scaled mean evolutionary rate of 6.6 x 10^−7^ substitutions per site per year for the whole genome, in line with estimates for other clonal human pathogens such as *Shigella sonnei* (6.0x10^−7^) and *Vibrio cholerae* (01 lineages; 8.0x10^−7^) (Holt et al., 2012; Mutreja et al., 2011).

Our divergence analyses for TPA samples provide a time to the most recent common ancestor (MRCA) less than 500 years ago (mean calendar year 1733, 95% HPD 1588-1848; Fig. 1B), which is after the start of the syphilis pandemic in the late 15^th^ century. Our findings point to the presence of a syphilis ancestor that originated at the dawn of the modern era and which succeeded in leaving a lasting genetic signature.

### Rapid spread of a contemporary epidemic cluster

A median-joining (MJ) network (Fig. 2A) for the 31 TPA samples highlights the mutational differences between haplotypes of the SS14 clade and the putative Nichols clade (both named after reference genomes) observed in the tree. The Nichols clade consists almost exclusively of samples collected from patients in North America from 1912 to 1986 and passaged in rabbits prior to sequencing, with the exception of one patient sample from 2013 (NE20). The clade contains a tight cluster of related sequences including the Nichols genome (Nichols-A), which have an MRCA in the 19^th^ century. In contrast, the SS14 clade has a more global distribution, encompassing European, North American and South American samples collected from infections between 1951 and 2013. Strikingly, the SS14 clade contains a dominant central haplotype (labelled as SS14-Ω in Fig. 2A) from which the other sequences radiate. The cluster associated with the SS14-Ω haplotype contains all but one of the recent patient samples from 2012-2013 (n=17) that were captured and sequenced directly, in addition to three samples from 1977 (n=1) and 2004 (n=2). Bayesian analyses estimate an origin for the SS14-Ω cluster in the second half of the 20^th^ century (median calendar year 1963, 95% HPD 1948-1974), at a time when incidence was reduced due to the introduction of antibiotics. The star-like topology of this cluster is suggestive of a recent and rapid expansion.

**Fig. 2.**
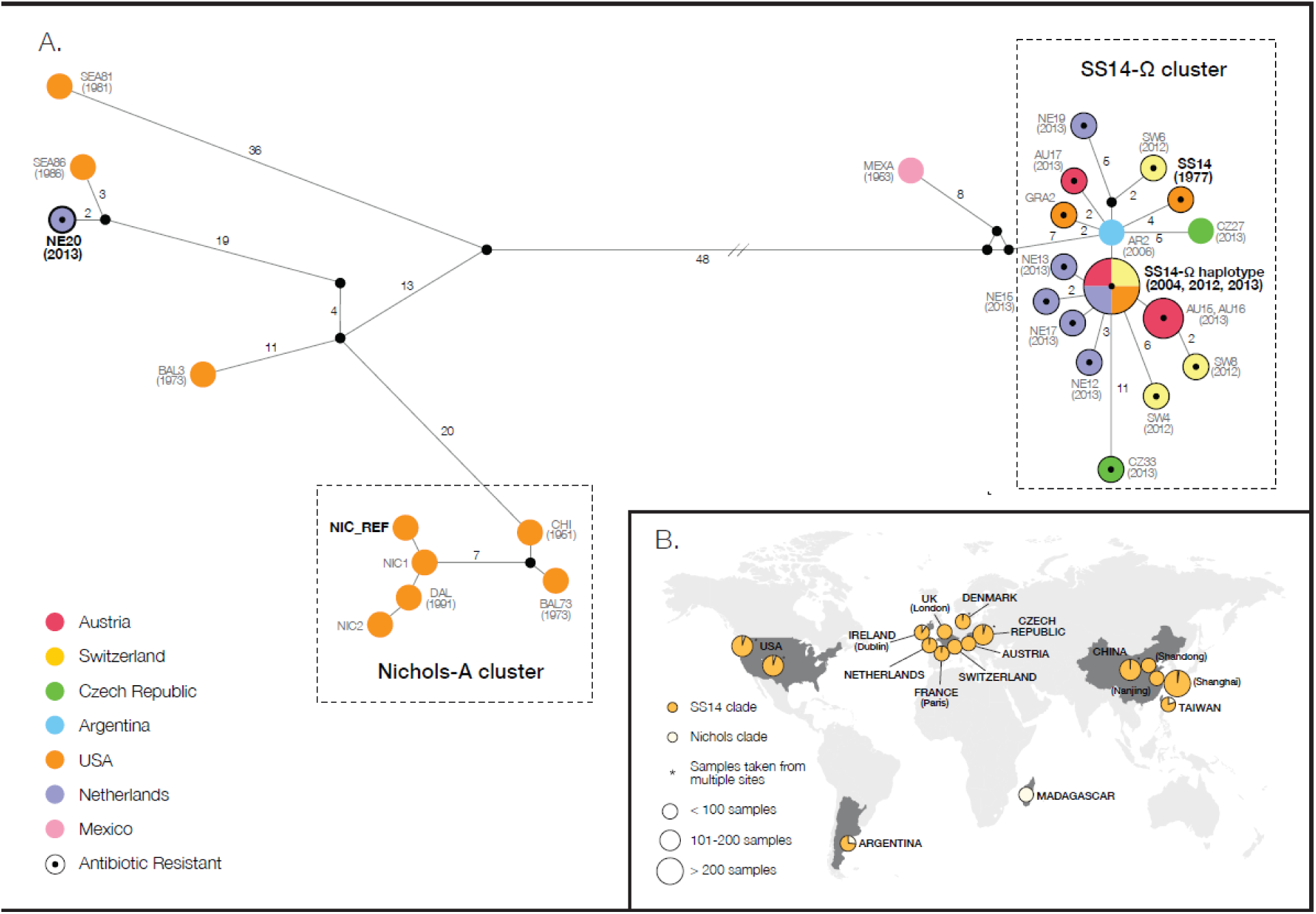
Median-joining (MJ) network analysis and geographic distribution of the SS14 and Nichols clades. A) Median-joining network for genome-wide variable positions after excluding sites with missing data (n=682). Circles represent haplotypes, with geographical origin color-coded. Number of mutations, when above one, is shown next to the lines. Inferred haplotypes (median vectors) are shown as black connecting circles. Central black circles within haplotypes indicate mutations associated with azithromycin resistance. B) Relative frequencies of SS14 versus Nichols clade isolates across the globe shown in the pie charts, with sizes proportional to sampling efforts. SS14 clade and Nichols classification are based on the TP0548 gene typing region (Christina M. Marra et al., 2010).

To determine whether the dominance of SS14 clade sequences applies across other countries for which genetic data is available, we examined the widely sequenced TP0548 hypervariable gene region from epidemiological typing studies (Christina M. Marra et al., 2010). This gene distinguishes the SS14 from the Nichols clade among TPA samples. Across 1353 worldwide TP0548 sequences, including the 78 from clinical samples in this study, we found that 94% of them fell into the SS14 clade, consistent with a probable recent spread of the epidemic cluster. The wide geographical distribution of the SS14 clade establishes it as representative of the present worldwide epidemic. While studies to date have focused on the Nichols strain (Lorenzo Giacani et al., 2012; Strouhal et al., 2007), our results indicate that further work on the SS14 clade is warranted.

Typing of samples collected across several different years in the Czech Republic, San Francisco, British Columbia and Seattle indicate that macrolide antibiotic resistance has increased over time (Grillová et al., 2014; C. M. Marra et al., 2006; Mitchell et al., 2006; Morshed & Jones, 2006; Stamm, 2010). We queried the presence of the two mutations (A2058G and A2059G) in the 23S rRNA genes associated with azithromycin resistance (Matejkova et al., 2009; Stamm, 2010). As observed in the MJ network, the resistance marker is a dominant characteristic of the SS14-Ω cluster (Fig 2A), although it is also found in the recent Dutch sample of the Nichols clade. Extending our analyses of the 23S rRNA gene to all sequenced samples from our study, including those with low coverage, revealed the mutations in 90% of the SS14 and 25% of the Nichols samples, indicating that neither resistance nor sensitivity is clade-specific. Hence resistance was not an ancestral characteristic of the SS14 clade. A likely scenario is that the extensive usage of azithromycin to treat syphilis and a wide range of bacterial infections, including co-infections with other sexually-transmitted diseases (STDs) such as chlamydia, has played an important role in the selection and subsequent spread of resistance (Geisler et al., 2015; Šmajs, Paštěková, & Grillová, 2015).

## Conclusions

We examined the genomic diversity of *Treponema pallidum* subsp. *pallidum* from syphilis samples isolated during the 20th and 21st centuries. The results present the first reported whole genome sequences successfully obtained directly from syphilis patients. Our analyses indicate that all TPA samples examined to date share a common ancestor that was infecting populations within the early centuries of the modern era. The present work does not necessarily resolve the question whether ancestral TPA originated in the Americas or Europe. However, it does suggest that events following the colonization of the Americas provided the context for the propagation of an STD lineage that has reached global proportions today. Furthermore, our analyses point to an epidemic cluster (SS14-Ω) that emerged after the discovery of antibiotics, and which displays the population genetic and epidemiological features of a pandemic.

## Acknowledgments

Research in Zurich was funded by the *Forschungskredit* and the University of Zurich. A.H was funded by the ERC Starting Grant APGREID. FGC and LSB were funded by project BFU2014-58565R from MINECO (Spanish Government). KIB was funded by Social Sciences and Humanities Research Council of Canada (756-2011-501). We are also grateful to Stephan Lautenschlager for his initial guidance. We thank Alexei Drummond for input on BEAST. We wish to thank Dr. Sheila Lukehart for providing HaitiB, Sea86-1, Bal3, Bal9, Bal73-1, and Grady1 strain DNA, and Dr. Christina Marra for providing UW249B and UW231B strain DNA. Special thanks to Lukas Keller and his lab for their support.

The copyright holders for this preprint are the authors. All rights reserved. No reuse allowed without permission.

